# Noncoding *de novo* mutations contribute to autism spectrum disorder via chromatin interactions

**DOI:** 10.1101/2019.12.15.877324

**Authors:** Il Bin Kim, Taeyeop Lee, Junehawk Lee, Jonghun Kim, Hyunseong Lee, Woo Kyeong Kim, Young Seok Ju, Yongseong Cho, Seok Jong Yu, Soon Ae Kim, Miae Oh, Tae Hwan Kwak, Sai Hali, Dong Wook Han, Eunjoon Kim, Jung Kyoon Choi, Hee Jeong Yoo, Jeong Ho Lee

## Abstract

Three-dimensional chromatin structures regulate gene expression across genome. The significance of *de novo* mutations (DNMs) affecting chromatin interactions in autism spectrum disorder (ASD) remains poorly understood. We generated 931 whole-genome sequences for Korean simplex families to detect DNMs and identified target genes dysregulated by noncoding DNMs via long-range chromatin interactions between regulatory elements. Notably, noncoding DNMs that affect chromatin interactions exhibited transcriptional dysregulation implicated in ASD risks. Correspondingly, target genes were significantly involved in histone modification, prenatal brain development, and pregnancy. Both noncoding and coding DNMs collectively contributed to low IQ in ASD. Indeed, noncoding DNMs resulted in alterations, via chromatin interactions, in target gene expression in primitive neural stem cells derived from human induced pluripotent stem cells from an ASD subject. The emerging neurodevelopmental genes, not previously implicated in ASD, include *CTNNA2*, *GRB10*, *IKZF1*, *PDE3B,* and *BACE1.* Our results were reproducible in 517 probands from MSSNG cohort. This work demonstrates that noncoding DNMs contribute to ASD via chromatin interactions.

---

For 276 ASD-affected Korean simplex families (Supplementary Table 1), we acquired average 69.12 *de novo* SNVs and 4.04 *de novo* INDELs per proband (and 67.83 and 4.02 per unaffected sibling), after controlling sample contaminations, inconsistent familial relationships, and coverage depth bias (see Methods) (Supplementary Fig. 1a-c). These numbers of *de novo* mutations (DNMs) were consistent with those reported from previous studies^1–3^. Validation rates were 95.4% (416 of 436) for *de novo* SNVs and 93.9% (46 of 49) for *de novo* INDELs, respectively. Our results confirm the increase in the *de novo* SNVs with parental ages^4, 5^ (Supplementary Fig. 1d,e). The mutation signatures of *de novo* SNVs converge on 1 (27%) and 5 (73%) (Supplementary Fig. 1f). This supported spontaneous mutagenesis in germlines as the underlying mutation mechanism, and previous studies reported that germline *de novo* SNVs best match signature 1 and 5^6, 7^. Taken together, the DNMs analyzed from our pipelines showed reliable mutation profiles including mean numbers and parental age correlations as well as mutation signatures compatible to previous large scale ASD whole-genome sequencing (WGS) studies.

In gene regulation, an enhancer interacts with a promoter via three-dimensional chromatin structures^8–10^. Exploiting the chromatin interactions between the DNase I hypersensitivity sites (DHSs) across the genome, we sought to identify target genes putatively dysregulated by enhancers carrying noncoding DNMs, which we defined as “NCD genes” (see Methods). We linked noncoding DNMs to NCD genes in each subject group: 230 to 363 in 276 Korean probands, 82 to 113 in 103 Korean unaffected siblings, and 450 to 632 in 517 MSSNG probands (Fig. 1a and Supplementary Fig. 2a). Mean numbers of the NCD genes per subject were slightly higher in the Korean and MSSNG probands than in the siblings (Korean probands=1.3, MSSNG probands=1.2, and Korean siblings=1.1), though there was no statistical significance of the differences (Supplementary Fig. 2b). The Korean and MSSNG probands had similar proportions of noncoding DNMs that are linked to multiple NCD genes (Korean probands=29%, MSSNG probands=28%, and Korean siblings=21%) (Supplementary Fig. 2c).

**Fig. 1.**
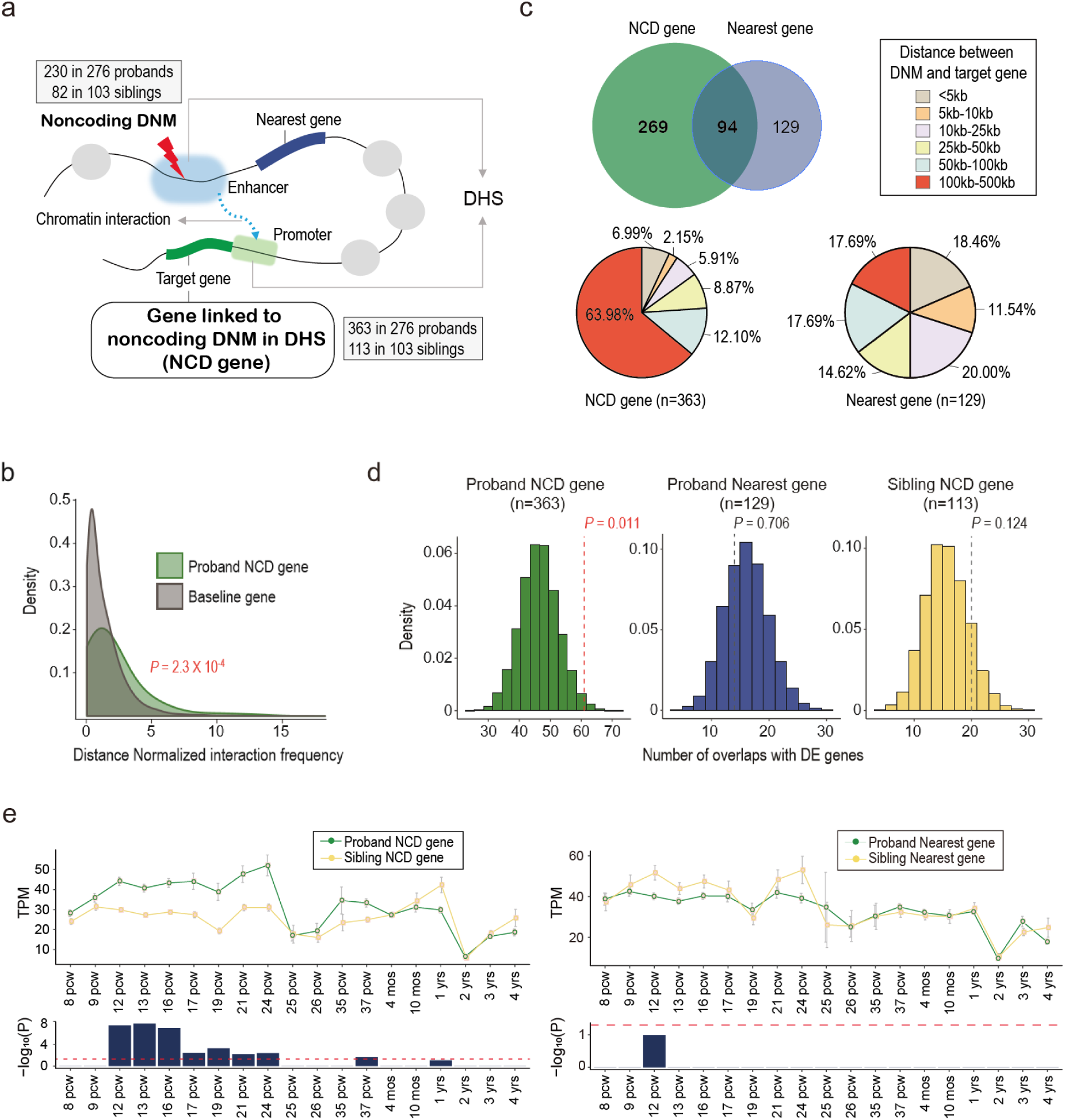
Target gene characterization using long-range chromatin interactions. **a**, Identification of NCD genes using chromatin interactions between distal enhancers and proximal promoters. **b,** Distribution of distance-normalized interaction frequencies for the NCD genes. *P* values were calculated using Student’s two-sided t-test and represent the significance of mean difference between interaction frequencies for the NCD genes and the baseline interaction frequencies. **c,** Comparison between the NCD genes and the genes nearest to noncoding DNMs. Among the 363 NCD genes, 269 (74.1%) were discordant to the genes nearest to the noncoding DNMs. Distances between DNMs and target genes were described for occupying proportions in the NCD genes and the nearest genes, respectively. **d,** Overlap with genes differentially expressed in ASD brains. Numbers of NCD genes in probands, nearest genes in probands, and NCD genes in siblings overlapping with the differentially expressed genes were simulated with 10,000 permutations, respectively. **e,** Expression patterns of the NCD genes and the nearest genes across brain developmental periods. The mean expression levels were compared between probands and siblings for the NCD genes and the nearest genes, respectively. *P* values were corrected for multiple tests with Bonferroni procedure. Error bars indicate standard errors. *TPM*, Transcripts per kilobase million; *PCW*, post-conception week; *mos*, months; *yrs*, years.

To validate those chromatin interactions within fetal human brains, we utilized published Hi-C data of neuronal progenitor cells^11^ as a baseline with which to compare Hi-C interactions for the NCD genes (see Methods). Reflecting the intensities of the Hi-C interactions, distance-normalized interaction frequencies for the proband NCD genes were significantly higher than those at baseline (Korean probands *P*=2.3×10^-4^, MSSNG probands *P*=5.0×10^-8^, Student’s two-sided t-test) (Fig. 1b and Supplementary Fig. 2d). These results indicated that the chromatin interactions between NCD genes and noncoding DNMs were validated using the independent Hi-C data.

Examining distances of the chromatin interactions, we observed, in the probands, 269 of the 363 NCD genes (74.1%) were discordant to genes nearest to the 230 DNMs (Fig. 1c). Moreover, 232 of the 363 NCD genes (63.9%) in the probands showed chromatin interactions over a distance of 100kb-500kb, whereas only 23 of the 129 genes nearest to the noncoding DNMs (17.8%) showed interactions over the same distance (*P*=2.4×10^-19^, Proportion test) (Fig. 1c and Supplementary Fig. 2e). These results suggested that majority of the noncoding DNMs might affect their target NCD genes via long-range chromatin interactions.

We then evaluated the functional properties of the 363 NCD genes in the probands in comparison to 129 of the nearest genes in the probands and to 113 NCD genes in the siblings. After 10,000 permutations, using genes differentially expressed in brains of ASD patients^12, 13^, we noted significant overlap only for the proband NCD genes (Proband NCD genes *P*=0.01, proband nearest genes *P*=0.71, and sibling NCD genes *P*=0.12, Random permutation test) (Fig. 1d). Besides, the expression levels of the NCD genes were significantly higher in the probands than the siblings during fetal periods of human brain development (Fig. 1e), implicating the fetal brain in the pathogenesis of ASD^14^.

Transcription factors act as key regulators of gene expression, and mutational changes in genomic sequences can disrupt the binding affinities thereof, altering transcription factor-dependent gene expression^15^. Accordingly, to delineate the transcriptional dysregulation by the noncoding DNMs, we analyzed their effect on binding affinities of transcription factors (see Methods). We focused on the noncoding DNMs affecting the NCD genes via chromatin interactions, which occupied only about 1.1% of all noncoding DNMs (Fig. 2a). The mean numbers of transcription factor binding sites disrupted by the noncoding DNMs affecting chromatin interactions were significantly higher in the probands than the siblings (Korean probands *P*=0.014, MSSNG probands *P*=0.002, Student’s two-sided t-test) (Fig. 2b). The probands showed greater number of DNMs that disrupted multiple transcription factor binding sites (≥4) than the siblings (*P*=0.01, Chi-squared test) (Supplementary Fig. 2f). Furthermore, we employed transcription factors from the Simons Foundation Autism Research Initiative (SFARI) that has curated ASD-risk genes, and examined the proportions of DNMs that disrupted corresponding transcription factor binding sites. Among the transcription factors implicated in the ASD-risk, majority showed greater enrichment of the disruptive DNMs in the Korean and MSSNG probands than the siblings (Korean probands=22 of 25 (88%), MSSNG probands=31 of 35 (88.6%), in comparison to Korean siblings) (Fig. 2c). In parallel, we also examined whether noncoding DNMs were implicated in transcriptional dysregulation regardless of their chromatin interactions with NCD genes. Among noncoding DNMs inside DHSs, we selected those without chromatin interactions that occupied about 40% of all noncoding DNMs (Fig. 2a). When we examined transcription factor binding site loss caused by the noncoding DNMs without considering chromatin interactions, no significant differences were observed between the probands and the siblings (Korean probands *P*=0.28, MSSNG probands *P*=0.66, Student’s two-sided t-test) (Fig. 2d,e). Taken together, these results suggested that the noncoding DNMs affecting NCD genes via chromatin interactions provoke transcriptional dysregulation implicated in ASD.

**Fig. 2.**
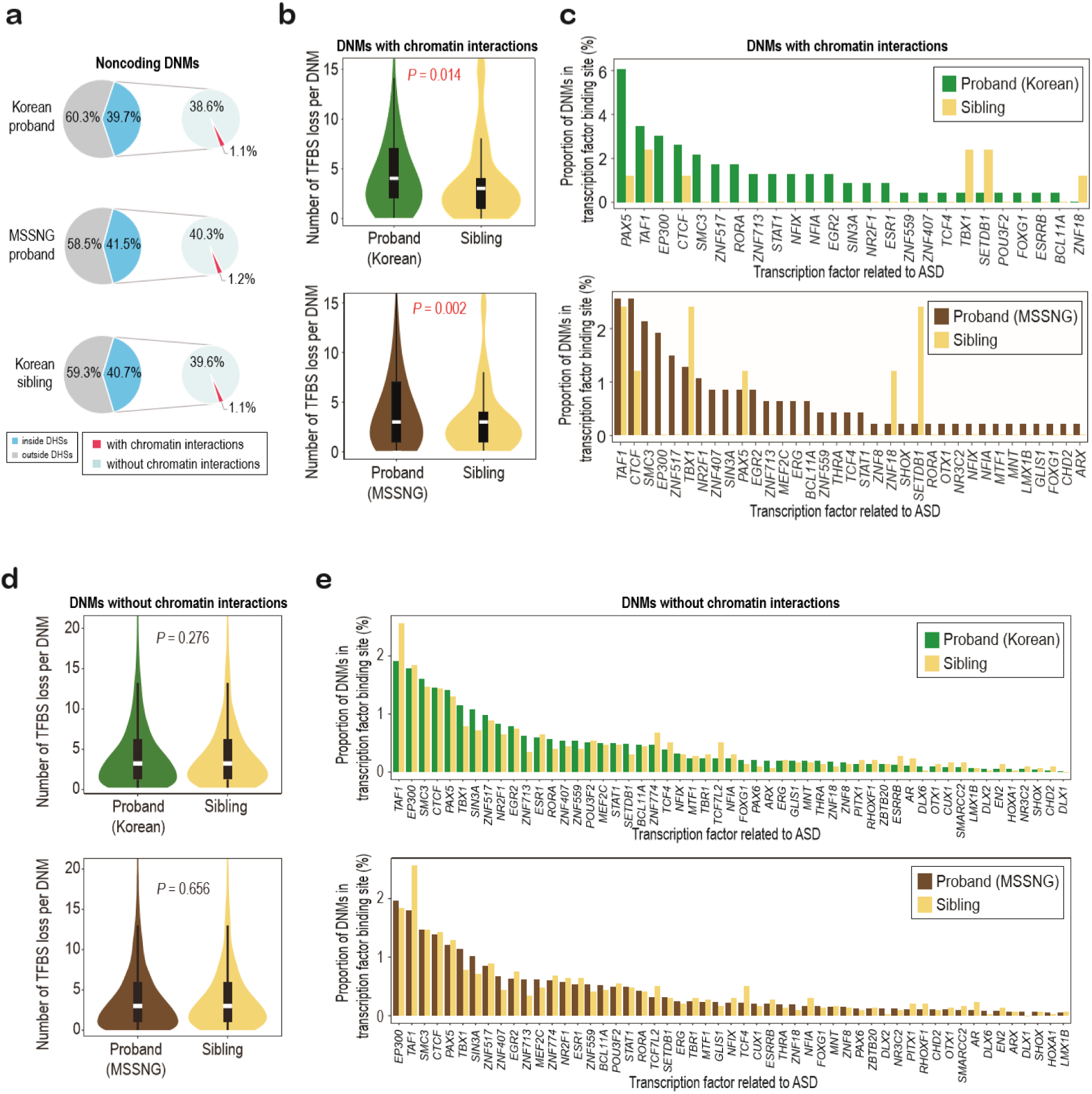
Transcriptional dysregulation by noncoding DNMs. **a**, Classification of DNMs according to overlapping with DNase hypersensitivity sites and their chromatin interactions. **b**,**d**, Number of binding sites lost due to DNMs with chromatin interactions (**b**) and without chromatin interactions (**d**). *P* values were calculated using Student’s two-sided t-test and represent the significance of differences in the numbers of transcription factor binding sites lost between probands and siblings. Boxplots depict the median and first and third quartiles; whiskers represent 1.5 times the interquartile range. **c**,**e,** Proportions of transcription factor binding sites lost due to DNMs with chromatin interactions (**c**) and without chromatin interactions (**e**).

To further emphasize the necessities of chromatin interactions in prioritizing the noncoding variants, we additionally sought to explore whether other methods not utilizing the chromatin interactions were able to detect any signals specific to noncoding DNMs in ASD probands compared to unaffected siblings^3, 16^. From a previous study^2^, we adopted its methodology in which numbers of the noncoding DNMs in DHSs were calculated in regards to any distances from ASD-risk genes. We compared the numbers of the noncoding DNMs between probands and siblings, and found no significant enrichment in the noncoding DNMs in probands compared to siblings (Supplementary Table 2). From another study^17^, we adopted disease impact score to prioritize meaningful noncoding variants in probands compared to siblings. In our dataset, the noncoding DNMs identified using chromatin interactions showed relative enrichment for high disease impact score (cut-off value=0) in probands compared to siblings, though this did not reach a statistical significance (Supplementary Table 3). This result seemed to be derived from the lack of annotation features relevant to three-dimensional chromatin structures in the disease impact score or from the size of Korean and MSSNG ASD cohorts (n=793). Therefore, these findings suggested that the chromatin interactions might be necessary for detecting functional noncoding variants in ASD.

Given genetic heterogeneity of ASD^18^, diverse mutation sources may affect genes crucial for neurodevelopmental disorders including ASD. We thus examined whether the noncoding DNMs affecting chromatin interactions recurrently dysregulated genes relevant to the neurodevelopmental disorders. We noted that 21 NCD genes were recurrently dysregulated by the noncoding DNMs via chromatin interactions in the Korean probands (Fig. 3a). Though none of the 21 NCD genes has been reported in the SFARI, some are associated with neurodevelopmental disorders: *DARS2* is related to leukoencephalopathy^19, 20^. *MASP1* is associated with learning disability^21^. *TAF2* is relevant to both microcephaly and mental retardation^22^. Additionally considering MSSNG probands, another 85 NCD genes were recurrently identified. Many of the identified NCD genes are also related to neurodevelopmental disorders including ASD (*ALOX5AP*^23^, *CACNA1C*^24^, *CNTN4*^25^, *MYO16*^26^, *SATB2*^27^, *SLC6A3*^28, 29^, and *XPO1*^30^), schizophrenia (*CACNA2D4*^31^, *CNTN4*^32^, *CPLX2*^33, 34^, *DLG1*^35^, and *MYO16*^36^), Down syndrome (*BACE2*^37, 38^), Joubert syndrome (*TMEM138*^39^), and attention deficit hyperactivity disorder (*CNOT10*^40^). These findings suggested that noncoding DNMs affecting chromatin interactions dysregulate converging risk genes giving rise to aberrant neurodevelopment.

**Fig. 3.**
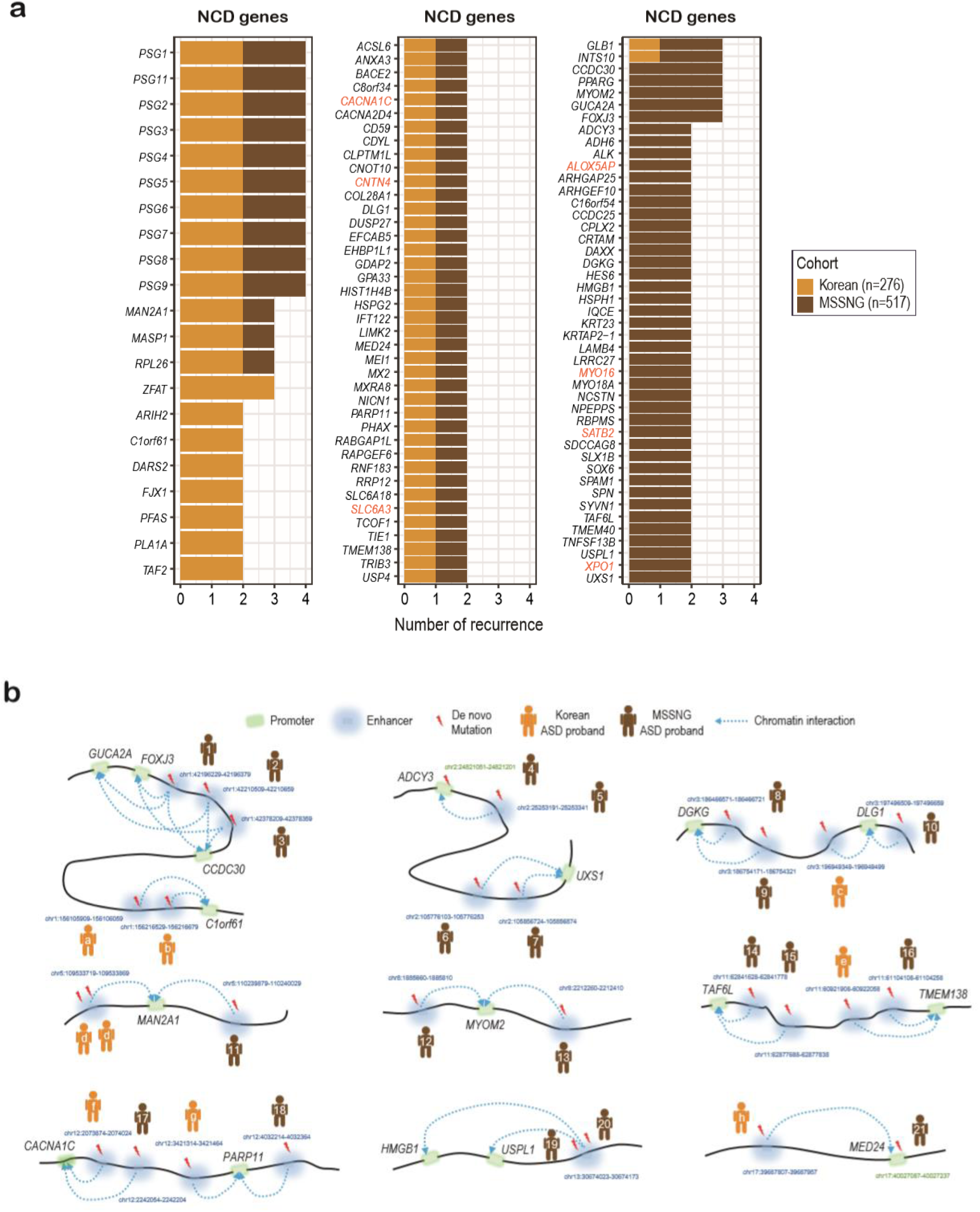
Genes recurrently dysregulated by noncoding DNMs via chromatin interactions. **a**, NCD genes that were recurrently dysregulated by the noncoding DNMs via chromatin interactions. Genes in orange indicate ASD-risk genes reported in Simons Foundation Autism Research Initiative. **b**, Visualization of chromatin interactions between NCD genes and noncoding DNMs. The examples depict chromatin interactions represented by both DNase-seq dataset (correlation coefficient≥0.8) and Hi-C dataset (distance-normalized interaction frequency≥2). Promoters (green), enhancers (blue), and chromatin interactions (blue dotted lines) are indicated.

For the recurrently identified NCD genes, we visualized some of the chromatin interactions linking the noncoding DNMs to the target genes (Fig. 3b). Those chromatin interactions were identified via DNase-seq dataset (Correlation coefficient ≥0.8) and validated using Hi-C dataset (Distance-normalized interaction frequency ≥2). Indeed, the NCD genes recurrently identified exhibited chromatin interactions of strong intensity. Furthermore, we tried to estimate the significance of the noted recurrences among the NCD genes. We found that calculating the probability of discovering recurrences among NCD genes revealed that this was more likely than would be expected if left to chance after 10,000 random permutations (*P*=0.0001 for recurrence ≥2, *P*=0.0007 for recurrence ≥3). Taken together, the noted recurrences among NCD genes as well as their biological relevance to neurodevelopmental disorders suggested that noncoding DNMs might be implicated in the ASD-risk via chromatin interactions.

The core phenotype of ASD may be represented by affected genes converging on specific networks and molecular pathogeneses^41^. We thus sought to examine whether NCD genes were expressed in ASD-associated brain regions, such as striatum^42^ and cerebellum^43^ during early brain development. To do so, we constructed 150 spatiotemporal expression signatures for individual genes using 15 brain regions and 10 developmental periods (see Methods). We found that NCD genes primarily overlapped with expression signatures in the fetal period (Cerebellum (*P*=0.007) and striatum (*P*=0.007) for NCD genes, Fisher’s exact test, Benjamini-Hochberg correction) (Fig. 4a). In parallel, in the probands, we also examined the expression signatures for genes putatively affected by damaging coding DNMs (Supplementary Fig. 3 and Supplementary Table 4,5) (see Methods), and found significant expression signatures of the genes across another ASD-related brain regions in the fetal periods (Sensorimotor cortex^44^ (*P*=0.0001) and amygdala^45^ (*P*=0.0005), Fisher’s exact test, Benjamini-Hochberg correction). Whereas, we found no significant expression signatures related to ASD for sibling NCD genes. These results suggested that noncoding DNMs contribute to the aberrant development of fetal brains via chromatin interactions, especially in collaboration with coding DNMs, in the ASD probands.

**Fig. 4.**
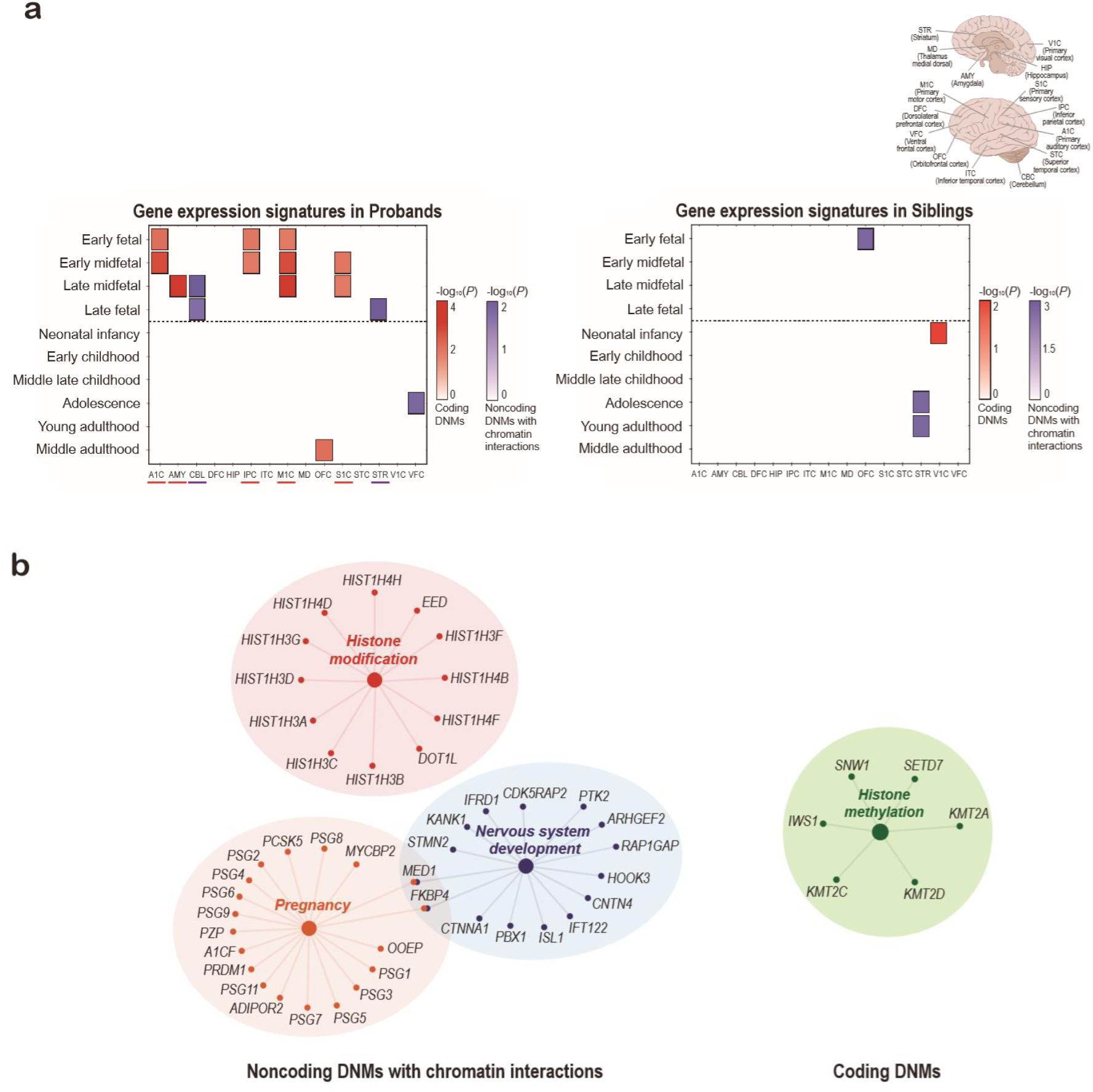
Biological convergence between noncoding variants with chromatin interactions and damaging coding variants. **a**, Spatiotemporal expression patterns of genes affected by noncoding and coding DNMs. Significance of enrichment in each of the 150 spatiotemporal expression signatures was calculated using Fisher’s exact test for NCD genes dysregulated by noncoding DNMs via chromatin interactions and for genes damaged by coding DNMs, respectively. Multiple *p* values were corrected using Benjamini-Hochberg procedure. **b**, Network clustering of genes affected by the noncoding and coding variants. The ClueGo package^69^ was utilized to depict clusters using Gene Ontology Biological Process and WikiPathways with a Bonferroni-corrected *p*-value threshold of 0.05.

We then explored which functional gene networks were affected by NCD genes, by analyzing Gene Ontology (GO) terms (Fig. 4b). We found that the terms including pregnancy and nervous system development were enriched for NCD genes, affirming a pathologic role of the noncoding DNMs with chromatin interactions in developing fetal tissues including brains (*P*=6.8×10^-19^ for pregnancy [GO:0007565], *P*=4.4×10^-12^ for regulation of nervous system development [GO:0051961], and *P*=2.0×10^-9^ for regulation of neuron differentiation [GO:0045665], Bonferroni correction). Histone modification was additionally enriched for NCD genes, implicating epigenetic dysregulation in the pathogenesis of ASD^46^ (*P*=4.0×10^-14^ for histone modifications [GO:0002369], Bonferroni correction). Of note, the genes affected by damaging coding DNMs were also found to be similarly enriched in terms related to the histone modification, (*P*=0.001 for regulation of histone methylation [GO:0031060], *P*=3.2×10^-4^ for histone lysine methylation [GO:0034968], and *P*=8.6×10^-4^ for histone H3-K4 methylation [GO:0051568]). This finding further suggested that there might be a collective pathogenesis between noncoding and coding DNMs that altogether contribute to the heterogeneous genetic architecture of ASD. Whereas, we found no significant terms implicated in the ASD-related pathogenesis for sibling NCD genes or for proband nearest genes. Taken together, these results indicated that noncoding DNMs contribute to the pathogenic mechanisms of ASD via chromatin interactions.

Noncoding DNMs that dysregulate NCD genes via chromatin interactions may represent a specific clinical feature in the probands. To examine that, we first classified probands by presence and types of the DNMs (Fig. 5a). As a result, we were able to detect 72.8% of the probands carrying either of the noncoding DNMs with chromatin interactions, damaging coding DNMs, or de novo CNVs. These proportions were similarly observed in the MSSNG probands and the unaffected siblings. We then focused on the 18.5% of the probands who had both the noncoding DNMs with chromatin interactions and the damaging coding DNMs, as we supposed that there would be a collective impact on any phenotypes of the probands. As expected, we found that the probands who had both the noncoding DNMs with chromatin interactions and the damaging coding DNMs showed lower mean intelligence quotient (IQ) scores than those with any one type of DNM and those with unexplored etiologies, such as inherited variants (Nominal *P*=0.017 and nominal *P*=0.031, respectively, Mann Whitney two-sided test) (Fig. 5b). Furthermore, probands with lower IQ scores tended to have larger numbers of both the NCD genes dysregulated by noncoding DNMs via chromatin interactions and the genes affected by damaging coding DNMs (Fig. 5c). The trend was not found for sibling NCD genes or for proband nearest genes to noncoding DNMs. Taken together, these results supported that multi-hits from diverse mutation sources including noncoding regions give rise to ASD with intellectual disability, and the NCD genes dysregulated by noncoding variants via chromatin interactions contribute to the oligogenic nature of the disorder^47^.

**Fig. 5.**
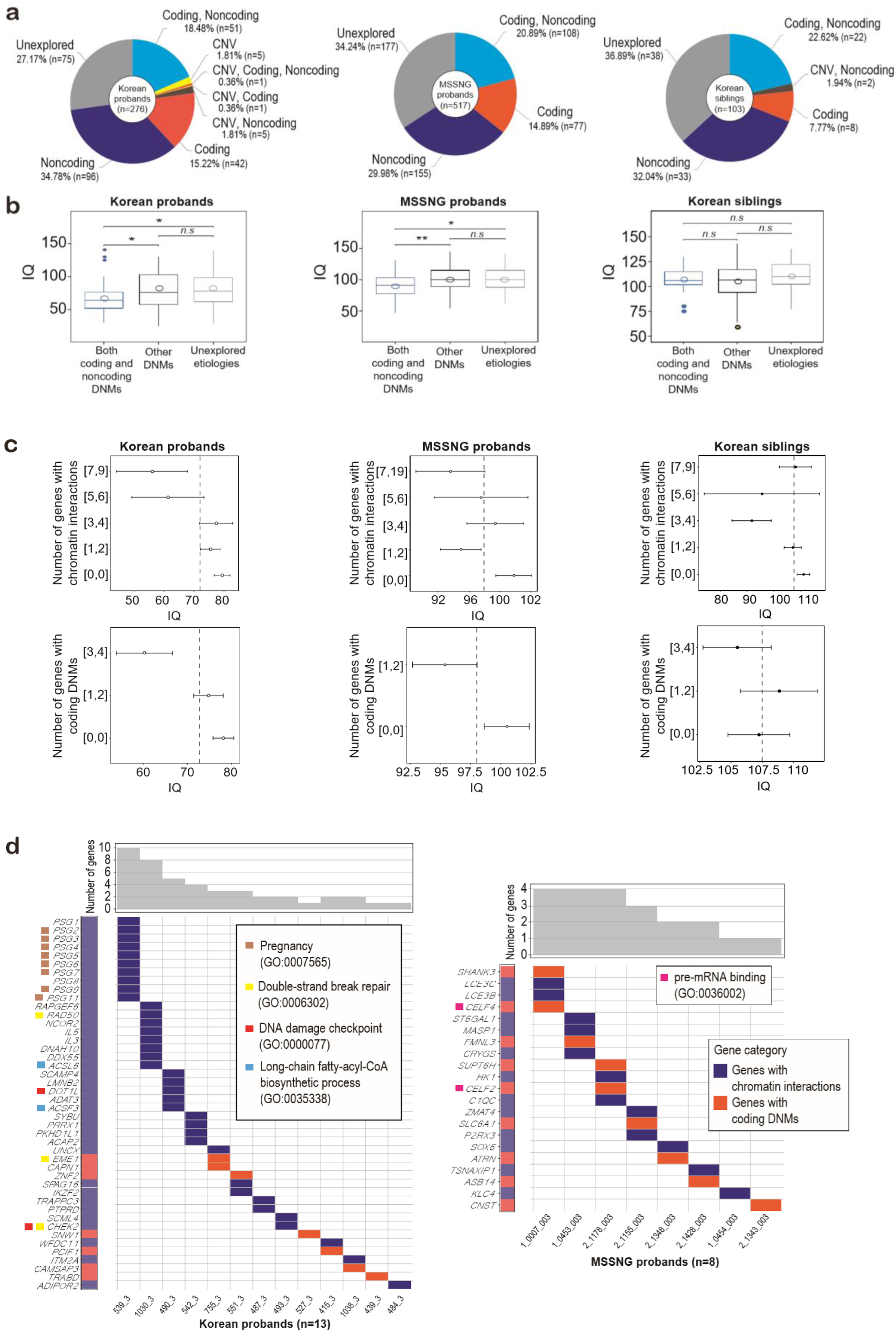
Collective impact of noncoding and coding variants on IQ in ASD. **a**, Classification of probands according to presence and types of DNMs. Probands with any of noncoding DNMs with chromatin interactions, damaging coding DNMs, or *de novo* copy number variations comprised 72.8% of all ASD probands. Of these, 18.5% carried both noncoding DNMs with chromatin interactions and damaging coding DNMs. **b**, Collective impact of the noncoding and coding DNMs on IQ in ASD probands. IQ scores were significantly lower in carriers of both noncoding and coding variants, compared to others. *P* values were calculated using Mann Whitney two-sided test. Hallow circles and horizontal lines within the boxplots indicate mean and median values, respectively. Error bars indicate standard errors. \**P*<0.05; \*\**P*<0.01; *n.s*, not significant. **c**, IQ scores and numbers of noncoding and coding variants. Probands with lower IQ scores tended to carry more genes dysregulated by the noncoding DNMs via chromatin interactions and more those affected by the damaging coding DNMs. Hallow circles indicate median values of the IQ scores in each bin of gene counts. Dotted lines and error bars indicate mean IQs and standard errors, respectively. **d**, General description of genes affected by noncoding and coding variants in probands with IQ scores in the lowest 10^th^ percentile. Gene Ontology terms were assessed for the affected genes in the selected ASD probands with intellectual disability, using EnrichR^70^ with a *p*-value threshold of 0.05.

We then further examined biological functions of the NCD genes dysregulated by noncoding DNMs via chromatin interactions and the genes affected by damaging coding DNMs in the probands with IQ scores in the lowest 10^th^ percentile (Fig. 5d). Among 45 genes in the selected Korean probands, we were able to extract significant GO terms including pregnancy [GO:0007565] (*P*=3.4×10^-12^), double-strand break repair [GO:0006302] (*P*=0.014), DNA damage checkpoint [GO:0000077] (*P*=0.04), and long-chain fatty-acyl-CoA biosynthetic process [GO:0035338] (*P*=0.028), potentially highlighting the biological background for low IQ in ASD. Taken together, these results indicated that the noncoding DNMs contribute to severe intellectual disability superimposed on ASD via chromatin interactions, in collaboration with damaging coding DNMs.

Lastly, to experimentally validate long-range disruptive effects of noncoding DNMs on the expression of NCD genes, we generated human induced pluripotent stem cells (hiPSCs) from a proband and his parents (Family ID 1289). The proband had four noncoding DNMs linked to eight NCD genes via chromatin interactions: chr7:50418513:C>T to *DDC*, *FIGNL1*, *IKZF1*, *GRB10*, and *ZPBP*; chr2:79656045:G>A to *CTNNA2*; chr11:14963517:G>C to *PDE3B*; and chr11:117170461:C>A to *BACE1*. Among established 18 hiPSC lines, we selected one line per donor based on morphology and transgene silencing (Fig. 6a and Supplementary Fig. 4a), and verified their pluripotency regarding typical marker expression and differentiation ability (Supplementary Fig. 4b-d). We induced differentiation of the hiPSCs into primitive neural stem cells (pNSCs), for which we confirmed the expression of typical markers of early neuroectoderms, such as PAX6, PLZF, and BLBP (Fig. 6b). By analyzing PAX6-positive cells, we assessed the differentiation efficiencies of the hiPSC lines and found them comparable with those of hESCs (Supplementary Fig. 4e). Gene expression analysis further revealed the activation of early neuroectodermal genes and silencing of pluripotency genes, indicating the successful differentiation of hiPSCs into a bona fide pNSC state (Fig. 6c).

**Fig. 6.**
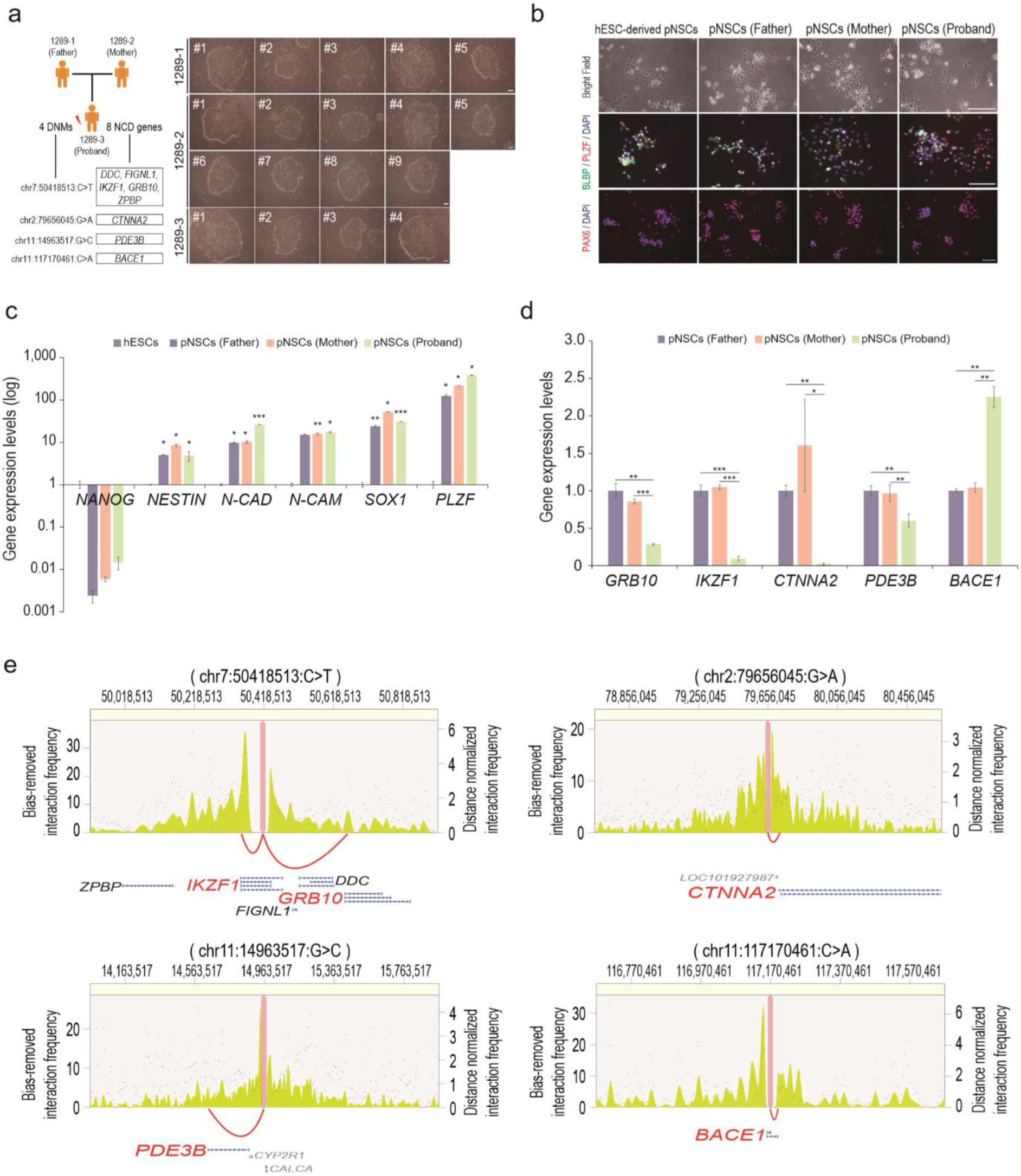
Transcriptional dysregulation of NCD genes in hiPSC-derived pNSCs with noncoding DNMs. **a**, Morphology of established hiPSC lines, as assessed by bright-field microscopy. Scale bars, 100 μm. **b**, The morphology of primitive neural stem cells (pNSCs) derived from hiPSCs was analyzed under bright-field (upper panel) and immunofluorescence microscopy. Antibodies against BLBP, PLZF (middle panel) and PAX6 (lower panel) were used. hESC-derived pNSCs were used as a positive control. Scale bar, 100 μm. **c**, Expressions of pNSC markers were analyzed by qPCR. Expression levels were normalized to those of hESCs. Error bars indicate the standard deviation (SD) of triplicate values. \**P*<0.05; \*\**P*<0.01; \*\*\**P*<0.001. **d**, Validation of target NCD genes in hiPSC-derived pNSCs by qPCR. Expression levels were normalized to those of pNSCs (ID 1289-1). Error bars indicate the SD of triplicate values. \**P*<0.05; \*\**P*<0.01; \*\*\**P*<0.001. **e**, Chromatin interactions between NCD genes and noncoding DNMs are described for four DNMs identified in the proband (ID 1289-3). Magenta dots and green peaks indicate bias-removed interaction frequencies (left axis) and distance-normalized interaction frequencies (right axis), respectively. Red arches indicate three-dimensional chromatin interactions at the neural progenitor cell stage. Genes in red indicate those with expression changes validated in this study. Variants at the top of the plots are equivalent to GRCh37.

Remarkably, at the pNSCs, the expression levels of five of the eight NCD genes identified in the proband were significantly different from those in his parents (Fig. 6d). In the proband, the expression levels of four NCD genes, including *GRB10*, *IKZF1, CTNNA2*, and *PDE3B,* were significantly lower, while *BACE1* expression was increased, compared to his parents, at the early neural stage. Interestingly, three of the validated NCD genes, including *GRB10*, *CTNNA2*, and *PDE3B*, were discordant to genes nearest to the noncoding DNMs, further supporting the importance of long-range chromatin interactions (Fig. 6e).

The NCD genes validated are of strong neurobiological interest. *CTNNA2* is primarily expressed in the developing cortex of mammals^48^. LOF mutations in *CTNNA2* in mouse models elicit defects in morphogenesis of dendritic spines^49^ and axonal projection^50^. A human iPSC model showed that bi-allelic loss of *CTNNA2* causes neuronal migration defects in the cerebral cortex^51^. *GRB10* is an imprinted gene at paternal alleles expressed in neurons^52^, and deletion of the paternal alleles has been found to hinder social behaviors in mice^53^. *GRB10* mediates a negative feedback loop in mTORC1 signaling^54^, which is linked to ASD^55^. Hyperactivity of mTORC1 alters the ratio of synaptic excitation to inhibition, causing ASD-like phenotypes in mice^56^. Additionally, *GRB10* is the top-ranked gene near the most down-regulated peaks of histone acetylation in postmortem brains of ASD patients^46^. *IKZF1*, known to facilitate differentiation of early cortical neurons, is highly expressed in mouse cortical progenitor cells at an early developing stage and gradually decreases in expression over time^57, 58^. *IKZF1* overexpression causes defective lamination during the late embryonic stage^58^, suggesting that time-wise regulation of its expression is important to proper neural development. In this context, low expression of *IKZF1* in the early neural stage may reflect erroneous neural development, although this warrants further study. *PDE3B* mediates hypothalamic leptin signaling (*PI3K*-*PDE3B*-cAMP), in which *PDE3B* is activated by leptin^59^. Leptin engages in hippocampal synaptic plasticity^60^, a mechanism linked to ASD^61^. In ASD patients, high leptin levels were frequently observed in blood^62^ and anterior cingulate gyrus tissue^63^, suggesting some form of leptin resistance in ASD^64^. Accordingly, reduced expression of *PDE3B* could reflect defective synaptic plasticity related to ASD. Future research, however, is required to obtain details on *PDE3B* and leptin-related synaptic plasticity. Unlike the four genes with reduced expression in the proband, *BACE1* showed increased expression in the pNSCs of the proband, compared to his parents. *BACE1* regulates neuronal proliferation^65^, and *BACE1* overexpression in mice induces microglia hyper-activation and aggressive synaptic pruning^66^. In keeping therewith, several researchers have argued that neuroinflammatory changes, such as over-pruning of synapses, could be pathogenic in ASD^67, 68^. Accordingly, the overexpression of *BACE1* in the early neural stage might reflect an aggressive neuroinflammation of ASD. Regarding the three NCD genes without any significant changes in their expression levels, we found that they had relatively low transcriptional activities and low H3K4me3 peaks, compared to the NCD genes with significant expression changes, in the early neural stages (Supplementary Fig. 4f,g). Taken together, these results indicated that the transcription of NCD genes in mutation-carrying pNSCs is significantly dysregulated via chromatin interactions, reflecting the contribution of mutations outside protein-encoding regions to the pathogenesis of ASD in early brain development.

In this study, to elucidate the genomic architecture underlying ASD, we generated and analyzed WGS data from parent-child simplex families in the Korean population (931 individuals). In parallel, we also analyzed WGS data from an independent dataset for ASD probands in the MSSNG database (517 individuals). In doing so, we identified noncoding DNMs that elicit transcriptional dysregulation of NCD genes via chromatin interactions in ASD. Furthermore, we found that the NCD genes dysregulated by noncoding DNMs contribute to ASD-relevant pathogenic mechanisms and low IQ, in collaboration with the genes affected by damaging coding DNMs. Overall, our findings suggested that variants outside protein-encoding regions confer major risks to ASD.

We specifically characterized the NCD genes putatively dysregulated by noncoding DNMs. Most (74%) of these genes were discordant to genes nearest to the noncoding DNMs, indicating the long-range chromatin interactions. Furthermore, numerous NCD genes were recurrently identified from the Korean and MSSNG probands. Many of these genes were related to neurodevelopmental disorders, which suggests that many mutational sources may converge on critical risk genes to elicit ASD. Particularly, to confirm the transcriptional dysregulation by noncoding DNMs, we validated expression changes in NCD genes in pNSCs differentiated from hiPSCs from a Korean ASD simplex family. The NCD genes validated, including *CTNNA2*, *BACE1*, *GRB10*, *IKZF1*, and *PDE3B*, were not previously spotlighted in the early neural pathogenesis of ASD. We suppose that those emerging genes can be novel ASD candidate risk genes, considering their transcriptomic changes in the pNSCs and literature-based crucial roles in the early neural development, although this warrants further studies. In conclusion, this study heightens our understanding of the heterogeneous genomic architecture of ASD, particularly target genes dysregulated by noncoding DNMs via chromatin interactions.

## Acknowledgement

This paper is dedicated to the memory of my colleague Seok Jong Yu, who died in 2019. We are grateful to all the families participating in this research, including Korean and MSSNG cohorts. This work was supported by grants from the Suh Kyungbae Foundation (to J.H.L.); Korean Health Technology R&D Project, Ministry of Health & Welfare, Republic of Korea (H15C3143 to J.H.L.); Brain Research Program through National Research Foundation of Korea (NRF) funded by the Ministry of Science and ICT (2017M3C7A1048092 to J.K.C.); National Research Foundation of Korea grant by the Ministry of Science and ICT (2017M3C7A1027467 to H.J.Y.); Research grant from Seoul National University Bundang Hospital (14–2015-011 to H.J.Y.); Institute for Basic Science (IBS) (IBS-R002-D1 to E.K.); Korea Institute of Science and Technology Information (K-19-L02-C07 to J.L., Y.C, and S.J.Y.)

## Author contributions

Experimental design, I.B.K., T.L., J.L., J.K., H.L., W.K.K., Y.S.J., Y.C., S.J.Y., S.A.K., M.O., T.H.K., S.H., D.W.H., E.K., J.K.C., H.J.Y., and J.H.L.; data generation, I.B.K., T.L., J.L., J.K., H.L., W.K.K., T.H.K., and S.H.; data processing. I.B.K., T.L., J.L., J.K., and H.L.; annotation of functional regions, I.B.K., T.L. and J.L.; data analysis, I.B.K., T.L., and J.L.; statistical analysis, I.B.K., T.L., and J.L.; manuscript preparation, I.B.K., T.L., J.L., J.K., J.K.C. and J.H.L.

## Data availability

All sequencing and phenotype data are hosted by the Korean Autism Genomics Database (KAGD) and are available for approved researchers at https:// kagd.kisti.re.kr.

## Code availability

Our codes for the main statistical analyses are uploaded on the KAGD, and are also available at GitHub. Any codes that are not uploaded are also available on request.

## Supplementary figures

**Supplementary Fig. 1.**
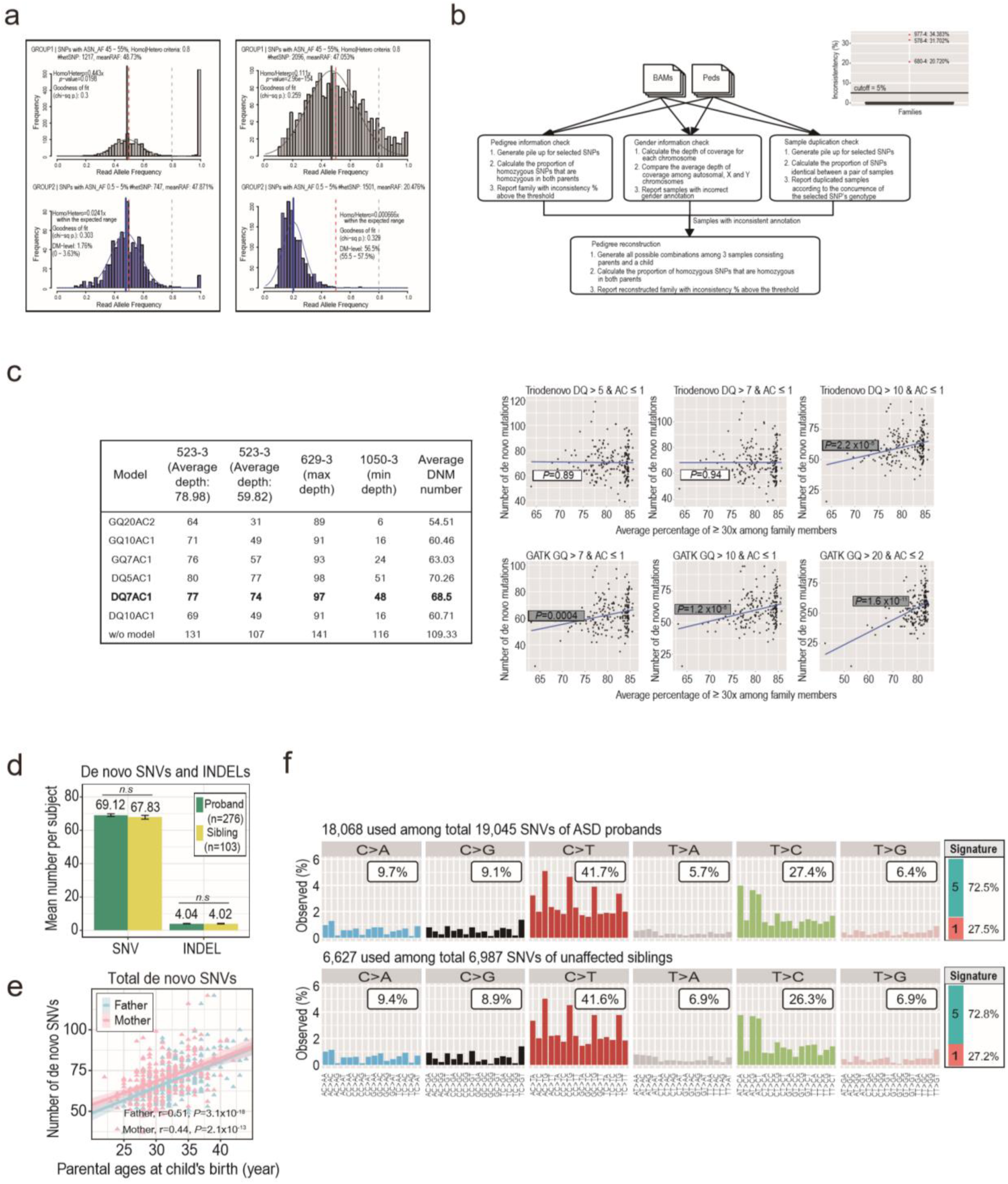
General profiles of DNMs identified in Korean ASD simplex families. **a**, Checking cross-contaminations between blood samples. Read allele frequencies (RAFs) of a sample were distributed across SNPs of Asian population at different allele frequency ranges (upper; 45-55% and lower; 0.5-5%). RAFs in each sample normally build two separate distributions of heterozygous and homozygous SNPs (Left), while RAFs in those with cross-contamination make shifted or irregular distributions (Right). **b**, Checking familial relationships among subjects. From bam files and ped files of subjects, we estimated the gender of each subject and the genotype concordance between parents and subjects by calculating the proportion of homozygous SNPs of a subject among the SNPs that are homozygous in both of parents (Left). Based on the genotype concordance proportion, we identified incorrectly annotated families that have more than 5% of inconsistent homozygous SNPs (Right). We further checked for the swapped samples and duplicated samples based on the genotype concordance proportions, and reconstructed the familial relationship among the subjects with incorrect annotations. **c**, Correction for coverage depth bias on the numbers of DNMs per subject. Average coverage depth among family members was significantly correlated with DNM number of a subject. Various calling models to correct the coverage depth bias on the DNM number were tested. Those tests resulted in our selection of the caller ‘TrioDeNovo’ that was applied subsequently to GATK HaplotypeCaller and Freebayes. **d**, Mean number of DNMs per subject. *P* values were calculated using Mann Whitney two-sided test. *n.s*, not significant. **e**, Correlations between DNM number and parental ages. There were strong correlations between the number of *de novo* SNVs and parental ages (Father r=0.51, *P*=3.1 x 10^-18^, Mother r=0.44, *P*=2.1 x 10^-13^, Pearson’s correlations) with an estimated increase of SNVs for each additional year of parental ages (Father 1.60 [1.27, 1.94], mother 1.44 [1.07, 1.80]). **f**, Mutation profiles and signatures. Signatures were estimated using linear regression option for maximum likelihood estimation, which is implemented in Mutalisk^71^. Cosine similarities indicating confidence for the proposed signatures (1 and 5) were 0.96 for probands and 0.95 for siblings, respectively.

**Supplementary Fig. 2.**
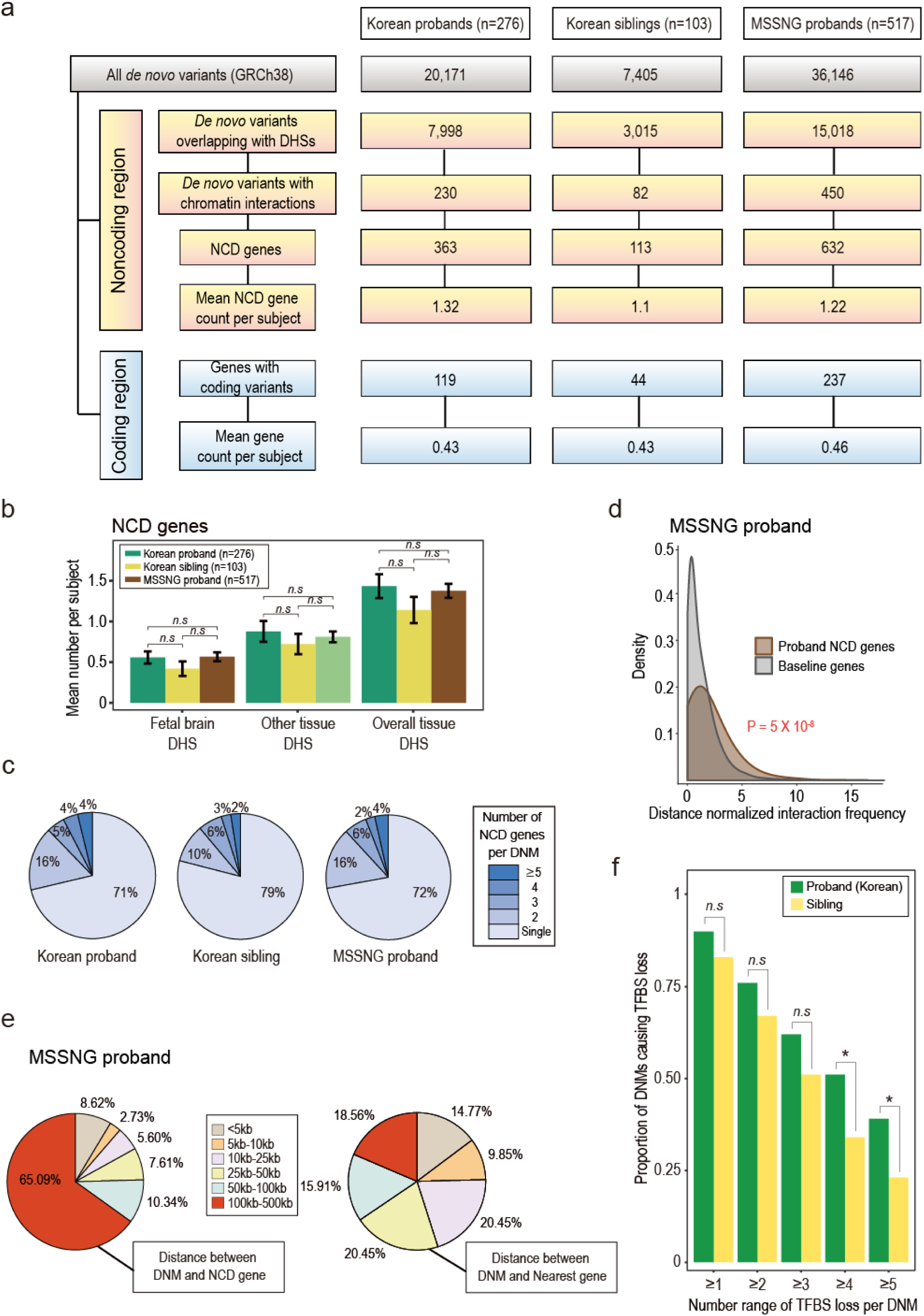
Profiles of noncoding DNMs and NCD genes coupled via chromatin interactions. **a**, Selection of noncoding DNMs and NCD genes based on chromatin interactions. NCD genes dysregulated by noncoding DNMs via chromatin interactions (yellow box) and genes affected by damaging coding DNMs (blue box) are respectively indicated for mean numbers per subject. MSSNG siblings were excluded in this study because of few subject numbers (n=42). *NCD genes*, target genes dysregulated by noncoding DNMs via chromatin interactions. **b**, Mean numbers of NCD genes per subject for fetal brain and other tissue DHSs. The mean numbers of Korean and MSSNG probands respectively were compared to that of Korean siblings using Mann Whitney two-sided test. *n.s*, not significant. **c**, Proportion of DNMs linked to multiple NCD genes. The Korean and MSSNG probands showed similar proportions of the DNMs linked to multiple NCD genes, compared to the Korean siblings (Korean proband=29%, MSSNG proband=28%, and Korean sibling=21%). **d**, Distribution of distance normalized interaction frequencies for NCD genes. *P* value was calculated using Student’s two-sided t-test and represents the significance of mean difference between the NCD genes’ Hi-C interaction frequencies and the baseline Hi-C interaction frequencies. **e**, Distance between noncoding DNMs and target genes. **f**, Proportions of DNMs causing multiple transcription factor binding site losses. The proportions of the DNMs causing the transcription factor binding site losses were compared between probands and siblings, using Chi-squared test. \**P*<0.05; *n.s*, not significant.

**Supplementary Fig. 3.**
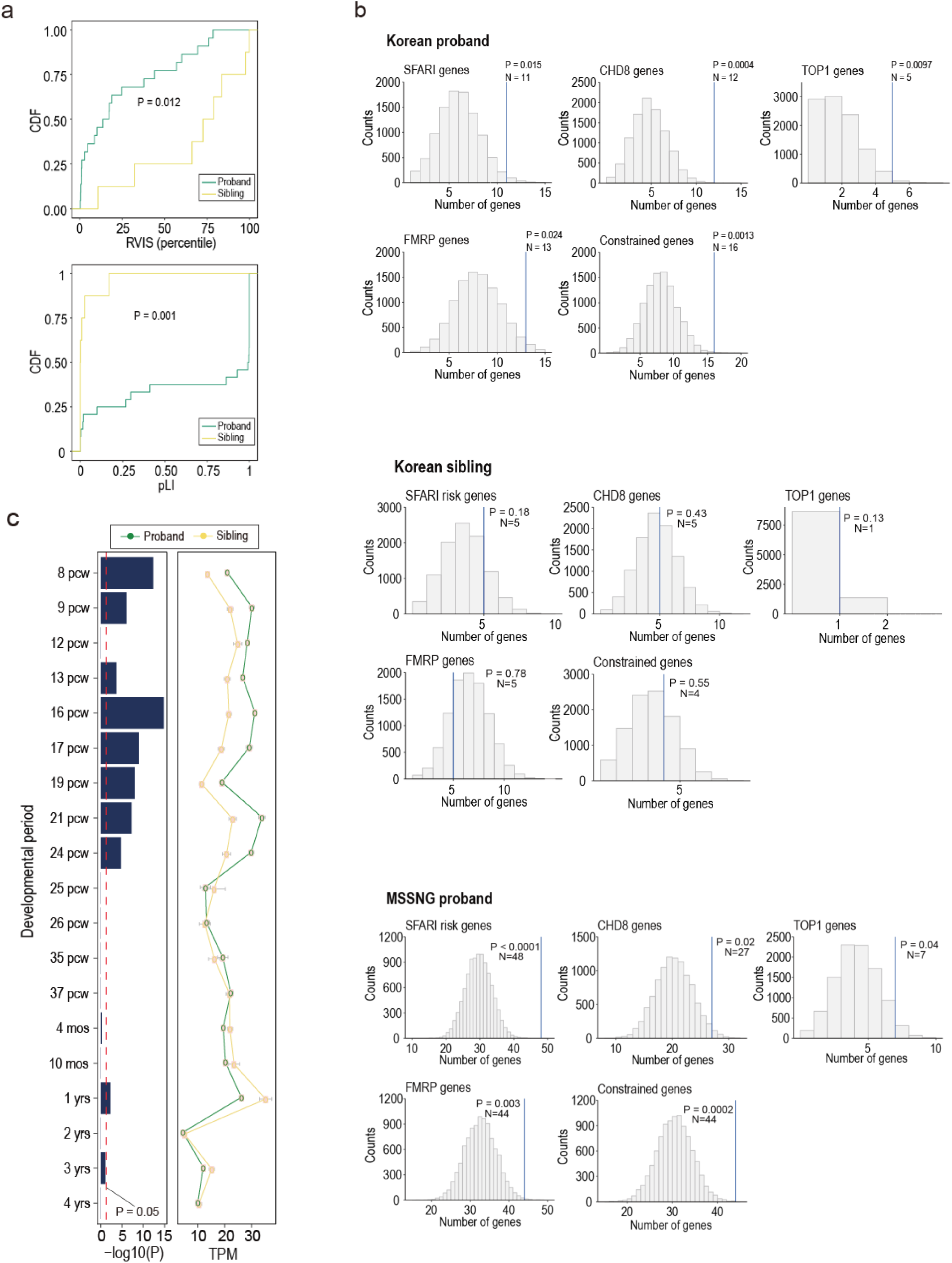
Functional characterization of genes affected by damaging coding DNMs. **a**, Cumulative distribution of mutation burden for damaging loss-of-function mutations. *P* values were calculated using Kolmogorov-Smirnov test and represent the significance of distribution differences for RVIS and pLI scores between probands and siblings. RVIS and pLI indicate residual variation intolerance score and probability of being loss-of-function mutation-intolerant, respectively. **b**, Enrichment test using published gene sets of ASD-risk and ASD-relevant pathologies (see Methods). *P* values represent statistical significance of observing the overlaps with the published gene sets, and were calculated by 10,000 permutations where genes were randomly extracted with re-sampling to reach the same numbers as the genes with coding DNMs identified from the cohorts. SFARI genes indicate ASD-risk genes classified as category 1 through 5, and syndromic. **c**, Expression patterns of genes with coding DNMs. *P* values represent the significance of mean differences for gene expression levels between probands and siblings, and were corrected for multiple tests with Bonferroni procedure. Mean expression levels of the genes were significantly higher in probands compared to siblings during the fetal period. Error bars indicate standard errors. *TPM*, Transcripts per kilobase million; *PCW*, post-conception week; *mos*, months; *yrs*, years.

**Supplementary Fig. 4.**
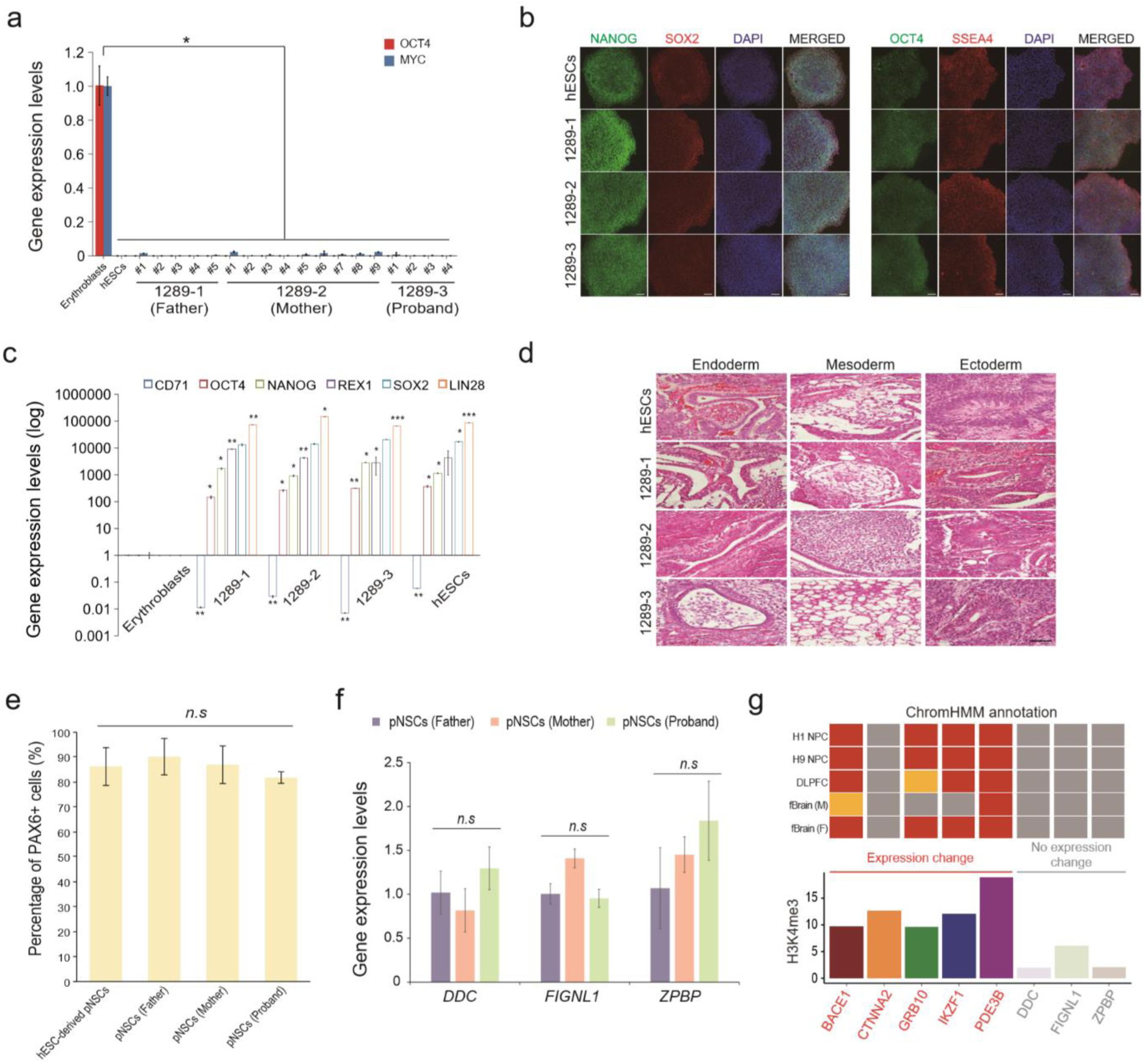
Generation and characterization of hiPSCs. **a**, The induction level of transgenes was analyzed by qPCR. Expression levels were normalized to those of erythroblasts transduced with reprogramming factors. Error bars indicate the standard deviation (SD) of triplicate values. \**P*<0.05. **b**, Immunofluorescence microscopy images of all hiPSC lines using antibodies against NANOG, SOX2, OCT4, and SSEA4. hESCs were used as a positive control. Scale bars, 100 *μ*m. **c**, Expression of pluripotency markers was analyzed by qPCR. Expression levels were normalized to those of erythroblasts. Error bars indicate the SD of triplicate values. *\*P<*0.05; *\*\*P<*0.01; *\*\*\*P<*0.001. **d**, Hematoxylin and eosin staining of teratomas derived from all hiPSC lines. Representative images display endoderm, ectoderm, and mesoderm-derived tissues. Scale bar, 100 *μ*m. **e**, The number of PAX6^+^ cells was counted at 2 weeks after neural induction. Error bars indicate the SD of triplicate values. *n.s,* not significant. **f**, Validation of the NCD genes in hiPSC-derived pNSCs by qPCR. Expression levels were normalized to those of pNSCs (ID 1289-1). Error bars indicate the SD of triplicate values. *n.s,* not significant. **g**, Transcriptional activities of the NCD genes. Compared to the three NCD genes with no expression changes, the five NCD genes with expression changes showed higher transcriptional activities in NPC, fetal brain and DLPFC (upper heatmap), and higher H3K4me3 peaks in NPC (lower barplot). In the heatmap, red, orange, and grey boxes respectively indicate transcription start site-related states, transcribed states, and repressed states, which were annotated by ChromHMM^72^. H3K4me3 peaks were independently adopted from 3D-genome Interaction Viewer and database^11^. *DLPFC*, dorsolateral prefrontal cortex; *fBrain*, fetal brain; *F*, female; *M*, male; and *NPC*, neural progenitor cell.

**Supplementary Table 1.** Sample phenotype summary.

|  | Overall |  | Males |  | Females |  |
| --- | --- | --- | --- | --- | --- | --- |
|  | ASD<br>(n=276) | Sibling<br>(n=103) | ASD<br>(n=238) | Sibling<br>(n=42) | ASD<br>(n=38) | Sibling<br>(n=61) |
|  | Mean (SD) | Mean (SD) | Mean (SD) | Mean (SD) | Mean (SD) | Mean (SD) |
| **Age at diagnosis (year)** | 7.4 (4.1) | 8.5 (4.9) | 7.6 (4.2) | 7.4 (4.4) | 6.7 (3.4) | 9.2 (5.1) |
| **WISC IQ** |  |  |  |  |  |  |
| Overall score | 76.7 (27.2) | 107.6 (17.4) | 78.2 (27.1) | 106.4 (21.4) | 67.6 (26.2) | 108.3 (14.5) |
| **Leiter IQ** |  |  |  |  |  |  |
| Overall score | 88.3 (30.8) | 112.8 (23.8) | 89.4 (31.1) | 108.7 (24.0) | 81.8 (29.3) | 115.6 (23.6) |
| **ADIR-A scale** |  |  |  |  |  |  |
| Overall score | 20.8 (5.9) | 2.5 (3.5) | 20.9 (6.0) | 3.4 (3.5) | 20.5 (5.6) | 1.9 (3.5) |
| **ADIR-B-verbal scale** |  |  |  |  |  |  |
| Overall score | 15.4 (4.6) | 1.7 (2.4) | 15.6 (4.6) | 2.5 (2.2) | 14.2 (4.3) | 1.3 (2.3) |
| **ADIR-B-nonverbal scale** |  |  |  |  |  |  |
| Overall score | 10.4 (3.1) | 1.1 (1.8) | 10.4 (3.1) | 1.5 (1.6) | 10.3 (3.1) | 0.8 (1.8) |
| **ADIR-C scale** |  |  |  |  |  |  |
| Overall score | 5.9 (2.5) | 0.5 (1.0) | 6.0 (2.5) | 0.9 (1.2) | 5.1 (2.2) | 0.3 (0.6) |
| **Social Responsiveness Scale** |  |  |  |  |  |  |
| Total t-score | 91.5 (27.5) | 23.5 (22.4) | 92.1 (27.6) | 28.3 (27.7) | 87.8 (26.6) | 20.3 (17.4) |
| **Social Communication Questionnaire** |  |  |  |  |  |  |
| Total score | 18.9 (8.1) | 3.7 (3.6) | 19.2 (8.1) | 4.1 (2.8) | 16.6 (7.7) | 3.4 (4.2) |

**Supplementary Table 2.**
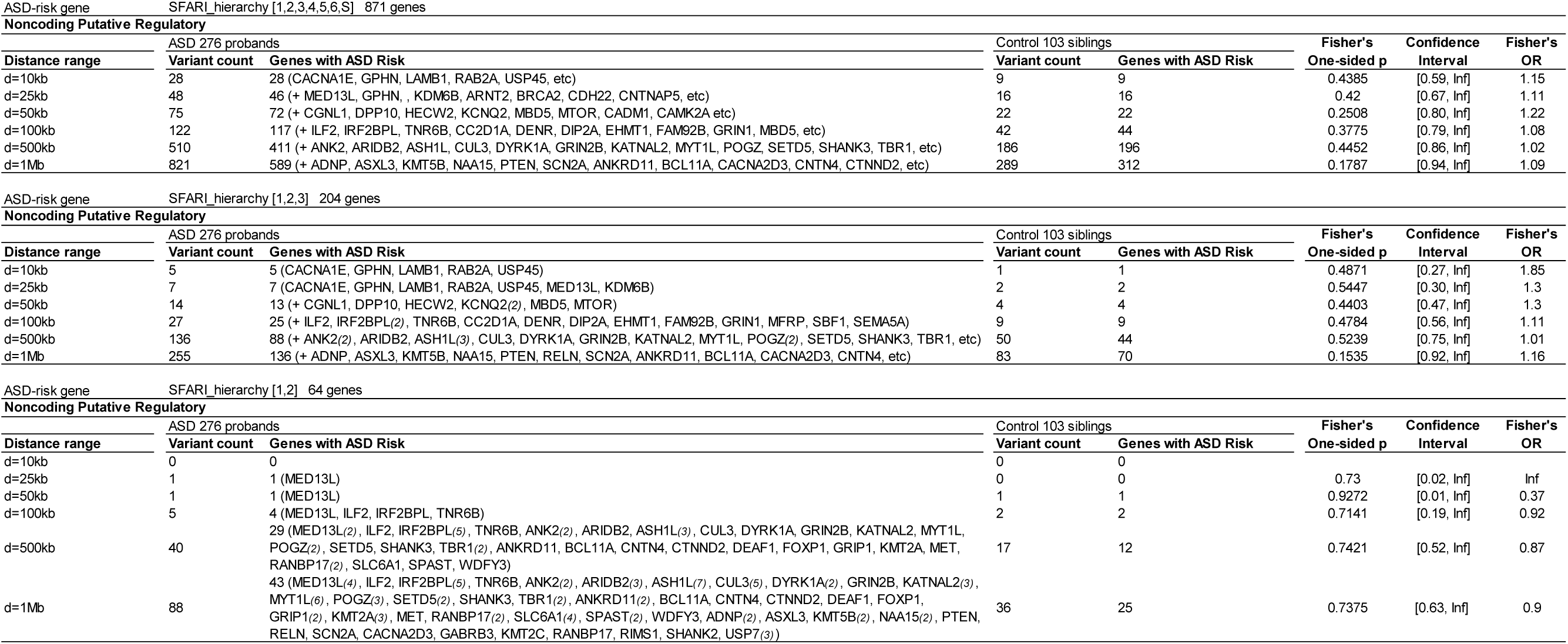
Proband and sibling comparison of counts of noncoding mutations in regulatory elements near ASD-risk genes.

**Supplementary Table 3.** Proband and sibling comparison of mutation counts for high disease impact score.

| Mutation class | Variants with DIS≤0 |  |  | Variants with DIS>0 |  |  | Odds Ratio | Confidence interval (95%) |  | Fisher's one-sided P-value |
| --- | --- | --- | --- | --- | --- | --- | --- | --- | --- | --- |
|  | ASD (n=793) | Sibling (n=103) | Sum | ASD (n=793) | Sibling (n=103) | Sum |  |  |  |  |
| Noncoding NCD | 381 | 48 | 429 | 302 | 31 | 333 | 1.227 | 0.744 | 2.047 | 0.472 |
| Noncoding (non-functional) | 34653 | 3374 | 38027 | 38967 | 3768 | 42735 | 1.007 | 0.959 | 1.057 | 0.785 |
| LOF | 62 | 7 | 69 | 20 | 2 | 22 | 1.129 | 0.193 | 11.986 | 1.000 |
| MIS | 213 | 25 | 238 | 60 | 5 | 65 | 1.408 | 0.500 | 4.908 | 0.642 |
| Synonymous | 149 | 22 | 171 | 40 | 6 | 46 | 0.984 | 0.355 | 3.171 | 1.000 |
| Coding (non-functional) | 273 | 29 | 302 | 45 | 14 | 59 | 0.341 | 0.160 | 0.758 | 0.007 |
| Overall counts | 35731 | 3505 | 39236 | 39434 | 3826 | 43260 | 1.011 | 0.963 | 1.061 | 0.659 |

**Supplementary Table 4.** Increased ASD risk with higher burden of impactful coding variants.

| Class | Korean probands (n=276) |  |  |  | Korean & MSSNG probands (n=793) |  |  |  | Korean unaffected siblings (n=103) |  |  |  |
| --- | --- | --- | --- | --- | --- | --- | --- | --- | --- | --- | --- | --- |
|  | Observed | Expected | Enrichment | P-value | Observed | Expected | Enrichment | P-value | Observed | Expected | Enrichment | P-value |
| MIS | 97 | 81.3 | 1.19 | 0.049 | 278 | 233.7 | 1.19 | 0.0026 | 34 | 30.4 | 1.12 | 0.28 |
| LOF | 24 | 11.3 | 2.13 | 0.0007 | 85 | 32.4 | 2.62 | $1.29 \times 10^{-14}$ | 10 | 4.2 | 2.37 | 0.01 |
| PROT | 121 | 92.6 | 1.31 | 0.003 | 363 | 266.1 | 1.36 | $1.04 \times 10^{-8}$ | 44 | 34.6 | 1.27 | 0.07 |

MIS, damaging missense mutations; LOF, damaging loss-of-function mutations; PROT, protein-altering mutations. PROT considers LOF and MIS simultaneously.

**Supplementary Table 5.** Genes with higher-than-expected mutation rates in ASD.

| Cohorts | Genes | LOF | PROT | LOF | PROT |
| --- | --- | --- | --- | --- | --- |
|  |  | Observed |  | Bonferroni-corrected *P*-value |  |
| Korean probands<br>(n=276) | *EMP1* | 1 | 1 | 0.01 | 0.15 |
|  | *OTOF* | 0 | 2 | 1 | 0.02 |
|  | *CHD8* | 1 | 2 | 0.23 | 0.02 |
|  | *SMAD6* | 1 | 1 | 0.03 | 0.54 |
|  | *PRPF18* | 1 | 1 | 0.03 | 0.33 |
|  | *ZNF362* | 1 | 1 | 0.04 | 0.49 |
|  | *ZNF2* | 1 | 1 | 0.04 | 0.50 |
| Korean & MSSNG<br>probands (n=793) | *SHANK3* | 2 | 3 | 0.001 | 0.003 |
|  | *CDK14* | 0 | 2 | 1 | 0.02 |

## Online Methods

### Sample collection

We collected 931 subjects from Korean simplex families with ASD probands to establish a comprehensive resource of genomic sequences and clinical phenotypes. Whole bloods were acquired from 552 parents, 276 probands and 103 unaffected siblings. DNA samples were extracted from the whole bloods according to QiaGEN producer’s protocols. We collected clinical information including ADI-R Diagnostic Algorithm Scores^73^, Wechsler IQ^74^, Leiter International Performance Scale IQ^75^, Social Responsiveness Scale (SRS)^76^, Social Communication Questionnaire (SCQ)^77^ as well as subjects’ age, sex, and sibling numbers. All phenotype information was cross-evaluated by two licensed psychiatrists. Whole bloods and clinical information were acquired with subjects’ agreement conformed to Institutional Review Board (IRB) protocols. IRB #: B-1606-350-303 (for whole genome sequencing), B-1609-361-004 (for organoid/human induced pluripotent stem cell repository), B-1607-353-004 (for organoid/human induced pluripotent stem cell analysis).

### Calling *de novo* SNVs, INDELs, and CNVs

Our purpose was to build accurate sequence data of Korean simplex families with ASD probands. Initially, fastq files were generated using Illumina Hiseq X at sequencing read depth 30x and mapped to the human reference genome (GRCh38) to produce BAM files using BWA-mem^78^. To find genuine calls, we used SNV and INDEL calls identified by both GATK HaplotypeCaller^79^ and FreeBayes^80^. Then we used TrioDenovo^81^ to call *de novo* SNVs and INDELs from the cohort and applied in-house filters adopted and modified from a previous study^82^ to acquire more specific *de novo* mutation calls. The followings are the filter conditions for *de novo* SNVs: 1) We removed SNVs near indels within +/- 5bp or low complexity regions, 2) SNVs in probands were removed if more than 70% of reads were called as heterozygous reference, or if more than 5% of non-reference reads occurred in either parent, 3) The call in the proband was considered false and removed if its sequencing depth was less than 10% of the total sequencing depth of the parents at the corresponding site, and 4) The ‘phred-scale’ filter based on likelihood of the genotype was further applied to refine the candidates (PL<10). Followings are the filters for *de novo* INDELs: 1) Depth for parents>10 and depth for probands>15, 2) QUAL>30, 3) QD>10, 4) The call was removed if it was reported in dbSNP, and 5) The call in probands was removed if more than 70% of reads were as heterozygous reference, or if more than 5% of non-reference reads occurred in either parent. The common filters for *de novo* SNVs and INDELs are as following: 1) We removed the call if the allele count was more than 2 in the multi-sample vcf, and 2) The call in the proband was removed if its allele frequency in East Asian populations of the 1000 genome project was more than 0.02%. To find accurate *de novo* CNVs, we applied Delly^83^, Manta^84^, and CNVnator^85^ to gain the reciprocally 30% intersected calls. Filter conditions were conformed to the CNVnator output forms and established as following: 1) Size≥1kb and <3Mb, 2) *p1*-value<0.05 and *p2*-value<0.05, and 3) *q0*<0.5. *De novo* CNVs were all confirmed via Integrative Genomic Viewer^86^. Sanger sequencing was applied to validate randomly extracted *de novo* SNVs and INDELs, and the validation rates of *de novo* SNVs and INDELs were 95.4% (416/436) and 93.9% (46/49), respectively.

### *De novo* calls of MSSNG probands

We sought to extract *de novo* SNVs and INDELs from the ASD probands of MSSNG cohort so that the DNMs could be comparable to those called in the Korean cohort. VCF files with *de novo* calls of probands were downloaded from MSSNG database (on September 17, 2017) (https://research.mss.ng). We set the filter conditions for selecting DNMs as following: 1) Relation: Proband, 2) Family type: Simplex, 3) Platform: Hiseq X or Hiseq 2000, 4) DNA source: Blood, and 5) Allele frequency (max) in 1000 genome project≤0.02%. Additionally, for autosomal chromosomes and female probands’ X chromosomes, we set as following: 1) Call depth≥10, 2) Reference allele count/call depth≤0.7, and 3) Inheritance: ‘0,0:0,0:0,1’ or ‘0,0:0,0:1,1’ or ‘0,0:0,0:1,2’. For male probands’ sex chromosomes, the conditions were as following: 1) Inheritance: ‘0,0:0,0:1,1’, 2) Call depth≥10, 3) Alternate allele count/call depth ≥0.8, and 4) GQ≥20. The DNMs of MSSNG probands were analyzed in parallel with those of Korean cohort to support the overall methodology used in this study.

### Correction for coverage depth bias on number of DNMs

DNAs extracted from whole bloods were sequenced with Illumina Hiseq X at different companies (Macrogen and Therragen in South Korea). We took into considerations, before sequencing, that different batches might be a bias on the *de novo* calls, and we provided the two companies with the identical blood samples without notice. We confirmed that batch effects of the different companies was not found. However, when we used GATK’s Genotype Refinement pipeline for determining *de novo* calls (with GQ>20, AC≤2), we identified that the numbers of DNMs were quite different between the two identical samples, which was biased by the average % of ≥30-fold coverage depth of the sample (1^st^ sequencing of subject ID 523-3, DNM number=64, 30x coverage depth(%)=78.98; 2^nd^ sequencing of subject ID 523-3, DNM number=31, 30x coverage depth(%)=59.82). We then found that the significant correlation between the average coverage depth among the sample’s family members and the number of DNMs per sample. Thus, we sought to find an algorithm of calling DNMs that could correct for the coverage depth affecting the number of DNMs. After applying several algorithms and cut-offs, we managed to conclude that the TrioDenovo^81^ with the conditions of DQ>7 and AC≤1 corrected for the bias on the number of DNMs.

### Defining NCD genes using chromatin interactions

To link noncoding DNMs to their target genes, we used DNase-seq dataset from a previous study^88^, where the strength of a chromatin interaction between a distal DHS (enhancer) and a proximal DHS (promoter) was characterized by Pearson correlation coefficients. We defined the significant correlation cutoff as r≥0.8 between the proximal and distal DHSs, with a distance between the two DHSs as 500kb apart at maximum. If a given DNM was located in the distal or proximal DHS of a gene, the gene was assigned as an ‘NCD gene’. To verify the chromatin interactions putatively affecting the NCD genes, we obtained Hi-C dataset of human neuronal progenitor cell from an independent study^11^. Distance normalized interaction frequency matrix in 5kb resolution was used to plot the baseline Hi-C distribution. The interaction frequencies between two bins containing the noncoding DNMs and the transcription start sites (TSSs) of the NCD genes were extracted from the baseline Hi-C distribution to separately plot the NCD genes’ Hi-C interaction distribution. Student’s two-sided t-test was used to assess the mean difference between the baseline Hi-C intensities and the NCD genes’ Hi-C intensities. To compare with the NCD genes, we extracted the nearest genes from the same noncoding DNMs that interact with the NCD genes, by using BEDtools^89^ and Gencode v28 TSS^90^.

### Differentially expressed genes

To compare the NCD genes with differentially expressed (DE) genes identified from independent patients with ASD, we collected the DE genes from two previous publications^12, 13^. In both studies, RNA-seq was previously performed for the brains of the patients and controls, respectively. The resultant DE gene set consisted of 2,666 genes, which was a union of the two resources. Random permutation test was performed to confirm whether the NCD genes of probands significantly overlapped with the DE gene set. Specifically, genes were randomly extracted from Gencode v28 protein coding genes^90^ to simulate the overlaps with the DE gene set. The number of overlaps with the DE gene set was inferred from the simulations, and *p* value was computed by repeating the permutation 10,000 times. The permutation tests were further performed for the nearest genes of probands and for the NCD genes of siblings, respectively.

### Transcription factor binding site loss

To identify the impact of DNMs on transcription factor binding sites (TFBSs), we utilized the position weight matrices in the TRANSFAC database^91^. The FIMO tool^92^ of the MEME SUITE package was used to determine the TFBS loss by DNM at *p*-value threshold of 1×10^-4^.

### Generation of hiPSC lines from PBMCs

Human peripheral blood mononuclear cells (PBMCs) were isolated by Ficoll-Paque Methods from whole blood^93^. Briefly, 8 ml of blood sample was diluted with PBS (1:1). The blood mixtures were transferred into 3 ml of Ficoll-Paque PLUS (GE Healthcare) and centrifugated for 30 min according to the manufacturer’s instructions. The PBMCs were carefully collected and transferred in a well of 12-well culture plate for expanding erythroblasts. 2×10^6^ PBMCs were cultured under expansion medium (EM) containing QBSF-60 (Quality Biologicals) supplemented with 50 ug/ml L-Ascorbic Acid (Biogems), 50 ng/ml SCF (R&D system), 10 ng/ml IL-3 (R&D system), 2 U/ml EPO (R&D system), 40 ng/ml IGF-1 (R&D system), and 1 uM Dexamethasone (Sigma). After expanding for 9-12 days, the cells were infected with lentiviral particles encoding hOCT4, hSOX2, hKLF4, and hMYC. The transduced cells were replated onto mouse embryonic fibroblast (MEF) feeder layer and maintained in E8 medium (Stem Cell Technologies). hiPSC colonies were formed at day 21 post-transduction. hiPSC colonies were manually picked and established as a cell line.

### Characterization of hiPSC lines

Total 18 hiPSC lines were generated from one ASD proband and his parents. The representative hiPSC line from each individual was selected based on their morphology and silencing of transgene expression. The representative hiPSC lines were used to evaluate their *in vitro* and *in vivo* pluripotency. hiPSCs were fixed with 4% paraformaldehyde and permeabilized with 0.3% Triton X-100/5% FBS in PBS for 2 hrs. The cells were incubated with primary antibodies for 16 hrs at 4°C. The appropriate fluorescent secondary antibodies were incubated for 2 hrs at room temperature. Nuclei were stained with Hoechst33342 (Invitrogen). Primary antibodies used for immunofluorescence are as follow: rabbit anti-NANOG (Cell Signaling, 1:500), goat anti-SOX2 (R&D system, 1:50), goat anti-OCT4 (Santacruz, 1:50), and mouse anti-SSEA4 (Merck, 1:100). qPCR was performed using SYBR Green PCR Master Mix (Applied Biosystems) on the ABI 7500 real-time PCR system (Applied Biosystems). Δ*Ct* values were calculated by subtracting the GAPDH Ct value from that of target genes. Relative expression levels were calculated using the 2^−ΔΔ^*Ct* Methods. Data are reported as mean values from triplicate measurements. Statistical significance was evaluated with unpaired two-tailed Student’s t-test.

For the *in vivo* differentiation assay (teratoma formation), approximately 2×10^6^ hiPSCs were mixed with Matrigel (Corning) and injected subcutaneously into the five-week-old immunodeficient NOD-SCID mice. After 6‒8 weeks, the teratomas were removed, fixed in 4% paraformaldehyde, and subjected to histological examination with hematoxylin and eosin staining.

### Differentiation of hiPSCs into neuronal cell lines

hiPSC lines were differentiated into primitive neural stem cells (pNSCs) by previously described protocol^94^. Briefly, the cells were cultured in the neural induction medium (NIM) containing DMEM/F12 (Corning) supplemented with 1x N2 (Gibco), 1x B27 without vitamin A (Gibco), 1% Penicillin/Streptomycin (Gibco), 1% GlutaMAX^TM^ (Gibco), 55 μM ß-mercaptoethanol (Gibco), 3 uM CHIR99021 (Tocris), 2 uM SB431542 (Peprotech), and 10 ng/ml hLIF (Millipore). After 7 days of neural induction, the cells were split 1:3 using TrypLE^TM^ (Gibco) for serial passages on the Matrigel-coated dish. After 2 or 3 passages, the neural rosette-like colonies were stably expanded. The expression levels of pNSC-related genes and the NCD genes were determined by qPCR. Statistical significance was evaluated with unpaired two-tailed Student’s t-test.

### Transcriptional activities of NCD genes

We sought to find biological backgrounds that possibly distinguish between the five NCD genes with the expression changes and the three without expression changes in the ASD family (ID 1289). To do that, we examined transcriptional activities of the 8 NCD genes in various cell types. We utilized ChromHMM where multiple chromatin datasets are integrated via machine learning, in order to predict chromatin state of a specific genomic region^72^. ChromHMM core 15-state model were collapsed into the following four categories: TSS-related states (TssA, TssAFlnk, TssBiv, BivFlnk), transcribed states (TxFlnk, Tx, TxWk), enhancer states (EnhG, Enh, EnhBiv), and repressed states (ZNP/Rpts, Het, ReprPC, ReprPCWk, Quies). We specifically assessed the TSS-related states and transcribed states of the 8 NCD genes in five cell types of Roadmap including H1 derived neuronal progenitor cultured cells (H1 NPCs), H9 derived neuronal progenitor cultured cells (H9 NPCs), male and female fetal brains (fBrains), and dorsolateral prefrontal cortex (DLPFC). Additionally, we employed 3D-genome Interaction Viewer and database (3DIV)^11^ to include otherwise missed transcriptional profiles of the NCD genes. H3K4me3 histone modification profiles were further examined in H1 derived neuronal progenitor cell from the 3DIV.

### Defining the coding DNMs putatively damaging genes

We used the Boruta package^95^ implemented in R to find putatively appropriate scoring tools for protein-coding variants that were likely to discriminate between probands and siblings. We were informed that the tools such as CADD^96^, Polyphen^97, 98^, PhyloP^99^, and PhastCons^100^ could be features discriminating between probands and siblings, while it warned GerpN, GerpNS and GerpS as unimportant features. We then built the in-house filter conditions for LOF and MIS as following. Here, LOF consists of frameshift, stop gained, start lost, splice donor and splice acceptor, while MIS includes in-frame deletion, in-frame insertion, and missense. For LOF mutations, either (1) CADD>20 and mammalian PhyloP>2.0, or (2) CADD>25 was applied. For MIS mutations, either (1) CADD>20, mammalian PhyloP>2.0, SIFT^101^<0.05, and Polyphen>0.45, or (2) CADD>20, primate PhastCons>0.9, mammalian PhastCons>0.9, and mammalian PhyloP>1.0 was applied. Those LOF or MIS mutations, based on the survival status under the filter conditions, were reclassified as damaging LOF, damaging MIS, and non-damaging LOF/MIS. We defined the genes with damaging LOF or damaging MIS as “the genes with damaging coding DNMs”. We equally applied the filter conditions to the coding DNMs of Korean probands, Korean siblings, and MSSNG probands.

### Gene burden implicated in ASD-related pathogenesis

We collected all genes with coding DNMs from Korean probands, Korean siblings, and MSSNG probands (471 genes, 118 genes, and 554 genes) to constitute each gene pool in each cohort. Then, we utilized reference gene sets of the ASD-risk^102^, the ASD-relevant pathologies such as FMRP (842 genes)^103^, CHD8 (1304 genes)^104^, and TOP1 (153 genes)^105^, and the high constraint (1003 genes)^106^. Here, we employed ASD-risk genes classified as category 1 through 5, and syndromic. To simulate the genes overlapping with each of the reference gene sets, we randomly selected 119 genes, 44 genes, and 235 genes (the same number of the genes with coding DNMs of each subject group) from the gene pools of Korean probands, Korean siblings, and MSSNG probands with 10,000 permutations (random re-sampling). From the permutation distributions for the counts of the genes overlapping with the reference gene sets, we estimated the significance by comparing the actual overlap counts with the distribution cutoff 5%.

### Coding DNM burden

We used denovolyzeR^107^ to test whether a subject group carries more coding DNMs than expected. We compared the expected number of DNMs with the observed numbers of LOF DNMs, MIS DNMs and protein-altering DNMs (LOF and MIS) from three groups: Korean probands, Korean & MSSNG probands, and Korean siblings. Additionally, we identified individual genes containing more LOF DNMs or protein-altering DNMs than expected by using the denovolyzeR.

### Developmental gene expression signature

The RNA-seq data of developmental gene expressions for human brains was downloaded from Brainspan^108^. Analysis was limited to 10 stages from early fetal (10–13 weeks postconception) to middle adulthood (up to 45 years) that included expression values (RPKM) for all 15 brain regions. We generated developmental signatures of gene expressions for each of the 150 combination windows from 10 stages and 15 regions, according to overall formulations of a previous method^109^.

Firstly, we calculated two z-scores for each gene i in terms of stage *s* and region *r*. Here, 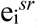 which is the gene’s expression in specific stage and region, was compared to the expression distributions across all stages at *r* to get 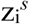 and across all regions at *s* to get 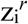 respectively.

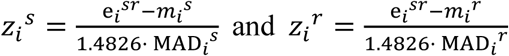

Where 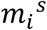 and 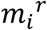 are the median expression levels of the gene i across all regions at the stage *s* and across all stages at the region *r*, and MAD*i^s^* and MAD*i^r^* are the median absolute deviations (MAD) of the gene expressions across all regions at the stage *s* and across all stages at the region *r*. These 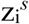 and 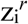 were then joined into a meta z-score.

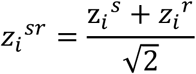

Finally, the genes with *Z_i_^sr^* ≥1.5 were utilized as the expression signatures for each of the 150 combination windows of stages and regions.

We then performed enrichment test of the 150 spatiotemporal expression signatures for the NCD genes dysregulated by noncoding DNMs via chromatin interactions and the genes affected by damaging coding DNMs, respectively. Fisher’s exact test was applied to calculate the significance of overlaps with all 150 expression signatures. *P* values were corrected for multiple tests using Benjamini-Hochberg procedure.

